# Detection of triglyceride metabolism in white blood cells using a novel fluorescently labeled long chain fatty acid analogue

**DOI:** 10.1101/2024.12.18.629134

**Authors:** Yasuhiro Hara, Ken-ichi Hirano

## Abstract

^123^I-15-(p-iodophenyl)-(*R*,*S*)-methyl pentadecanoic acid (^123^I-BMIPP) is a long-chain fatty acid (LCFA) analog developed to examine myocardial LCFA metabolism and has been used as a tracer for nuclear cardiology. However, its use is limited because of the specialized features of cardiac scintigraphy. In this study, a novel BMIPP-based probe, in which iodine-123 was replaced with a fluorescent compound, was used to extend the application of ^123^I-BMIPP to *ex vivo* analysis of fatty acid metabolism. To confirm that fluorescently labeled BMIPP-based probe (fluorescent BMPP) is effective for the detection of LCFA metabolism *ex vivo*, we performed fluorescence-activated cell sorting (FACS) analysis for the incorporation of fluorescent BMPP using white blood cells from human peripheral blood. FACS analysis showed that the incorporation of fluorescent BMPP into white blood cells was dose-dependent. Fluorescence intensities of cells incorporating fluorescent BMPP were attenuated after incubation in non-fluorescent BMPP medium, suggesting that these cells could export the probe. Neutrophils from a patient with primary triglyceride deposit cardiomyovasculopathy, a rare type of triglyceride (TG) metabolism disorders, showed lower BMPP export and was ameliorated after treatment with tricaprin. These results suggest that fluorescent BMPP could be imported to and exported from cells through the same mechanism as ^123^I-BMIPP, reflecting the mechanism that occurs for LCFAs during TG metabolism. This study shows that fluorescent BMPP has the potential to be used as a probe for the diagnosis of TG metabolic diseases *ex vivo*.

## 1. Introduction

Our goal is to develop a useful *ex vivo* diagnostic system that can examine intracellular triglyceride (TG) hydrolysis in different types of heart disease using peripheral white blood cells (WBCs). WBCs use fatty acids (FAs) as an energy source when they are circulating and non-activated [1,2], similar to the heart. One of our target diseases is primary triglyceride deposit cardiomyovasculopathy (TGCV). This disease is caused by a genetic deficiency of adipose triglyceride lipase (ATGL), a rate-limiting enzyme for intracellular TG hydrolysis, resulting in excessive accumulation of TG in cardiomyocytes and the coronary artery [3-5]. We previously established diagnostic criteria for TGCV and demonstrated the increasing importance of ^123^I-15- (p-iodophenyl)-(*R*,*S*)-methyl pentadecanoic acid (^123^I-BMIPP) cardiac scintigraphy using single-photon emission computed tomography (SPECT) imaging [5-7]. The long-chain FA (LCFA) analog, ^123^I-BMIPP, has been used in nuclear cardiology to study FA metabolism in different types of heart disease [8-11], including TGCV. Slow oxidation receptivity attributed to the methyl group at the β-3 position allows ^123^I-BMIPP to be a powerful probe for detecting FA flux abnormalities in patients’ hearts.

However, the use of iodine-123 and scintigraphy with SPECT equipment is associated with several disadvantages, including increased patient burden and the need for appropriate facilities. Therefore, we focused on developing a novel fluorescently labeled BMIPP-based structure where the radioisotope, iodine-123, was replaced with a fluorescent compound and could act as an LCFA analog [12]. In the present study, we investigated the potential of fluorescent BMPP for *ex vivo* analysis of LCFA metabolism using WBCs. We demonstrated that fluorescent BMPP was incorporated into cells in a concentration-dependent manner and exported from cells after incubation with fluorescent BMPP-deprived medium. We confirmed that cells derived from patients with primary TGCV showed decreased export of fluorescent BMPP and was ameliorated after treatment with tricaprin.

## 2. Materials and methods

### 2.1 Treatment of whole blood cells with Alexa680-BMPP

Alexa680-BMPP was prepared as described in a previous report [12]. Samples of 3 mL fresh peripheral blood from healthy volunteers or patients with a CD34 deficiency were collected in heparinized tubes. A 10× concentration of Alexa680-BMPP was prepared in phosphate-buffered saline (PBS) and added to 200 μL aliquots of whole blood in microtubes to produce the desired concentrations, as indicated in Figure (0.2 × 3^-3^ to 0.2 × 3^0^ μM). The samples were incubated for 30 min at room temperature, followed by the addition of 2 mL BD FACS lysing solution (BD Biosciences, San Jose, CA) to lyse erythrocytes that were then removed by centrifugation (120 × *g* for 10 min). The remaining WBCs were resuspended in 600 μL PBS followed by FACS analysis.

### 2.2 FACS analysis of Alexa680-BMPP incorporated into WBCs

A MacsQuant Analyzer flow cytometer (Miltenyi Biotec, Bergisch Gladbach, Germany) was used to measure side scatter (SSC) and forward scatter (FSC) light and FL-6 channel (excitation and emission at 635 nm and 655–730 nm, respectively) for Alexa680 fluorescence according to the manufacturer’s instructions. Before measuring the Alexa680-BMPP-treated samples, SSC and FSC amplifications were adjusted using cells obtained in the same protocol immediately after adding Alexa680-BMPP to preliminarily determine gating regions containing the major leukocyte populations (lymphocytes, monocytes, and granulocytes). The Alexa680 fluorescence parameter was adjusted using these cells to determine the appropriate peak positions. Then, 10^4^ cells in 200–400 μL PBS were measured for the one-color fluorescent analysis of Alexa680. Additionally, the FL-6 channel (excitation and emission at 488 nm and 525 nm, respectively) was used to measure the fluorescence intensity of fluorescein. Fluorescein-labelled BMPP was kindly provided by Dr. Kenji Monde (Hokkaido University, Japan).

### 2.3 Preparation of white blood cell (WBC) fractions

The WBC fraction was obtained from whole blood by removal of erythrocytes using Hetasep (StemCell Technologies, Vancouver, Canada). Briefly, 6 mL fresh peripheral blood from healthy volunteers or a patient with primary TGCV was incubated with a 5:1 ratio of Hetasep for 30 min at 37 °C to remove erythrocytes by gravity sedimentation. WBCs in the upper layer were collected and washed twice with Dulbecco’s Modified Eagle Medium (DMEM) using gentle centrifugation (120 × *g* for 10 min), and then resuspended in 2.2 mL DMEM followed by the Alexa680-BMPP-incorporation and export assay.

### 2.4 Alexa680-BMPP-incorporation and export assay using WBC fractions

A 10× concentration of Alexa680-BMPP in PBS was prepared and added to 400 μL aliquots of the WBC fractions in microtubes to produce a final concentration of 0.075 μM. The assay was performed as follows: after 30 min incubation at room temperature for Alexa680-BMPP incorporation, the samples were immediately added to 4 mL BD FACS lysing solution to stop the reaction and remove erythrocytes completely. The samples were then analyzed using FACS as described in section 2.2. On the other hand, the export assay was performed using the remaining incorporation samples as follows: the samples were washed twice with fresh DMEM using centrifugation (120 × *g* for 5 min) to remove Alexa680-BMPP from the extracellular fluid, resuspended in 500 μL fresh DMEM, and then incubated at 37 °C for different time periods, as indicated in Figure (10 to 120 min). After incubation, the samples were added to 5 mL BD FACS lysing solution and subjected to FACS analysis.

## 3. Results

### 3.1 White blood cells derived from human peripheral blood incorporated Alexa680-BMPP

In this study, we used the Alexa680-labeled BMIPP basic skeleton (Alexa680-BMPP) described in our previous report [12]. To evaluate the possibility of using Alexa680-BMPP for *ex vivo* analysis of WBCs, we employed a simple method in which Alexa680-BMPP was directly added to peripheral blood samples. After Alexa680-BMPP treatment and the lysis and removal of erythrocytes, the fluorescence intensities of cells corresponding to neutrophils, monocytes, and lymphocytes were selectively analyzed using FACS (Fig. 1A). FACS analysis showed that the fluorescence intensities of each cell population markedly increased depending on the concentration of Alexa680-BMPP, whereas those of the Alexa Fluor 680 control only slightly increased when higher concentrations were used (Fig. 1B). The slight increase in the fluorescence intensity of the Alexa Fluor 680 control might have occurred via simple passive diffusion. Thus, these results confirmed the dose-dependent incorporation of Alexa680-BMPP into cells as an LCFA analog. To eliminate the possibility of non-specific import through specific characteristics of the Alexa680 structure, we examined the incorporation of fluorescein-labeled BMPP using FACS and confirmed that it was also incorporated in a dose-dependent manner (Fig. 1C). These results suggest that fluorescently labeled BMPP could be imported into WBCs via a transport pathway similar to that of LCFA.

**Figure 1.**
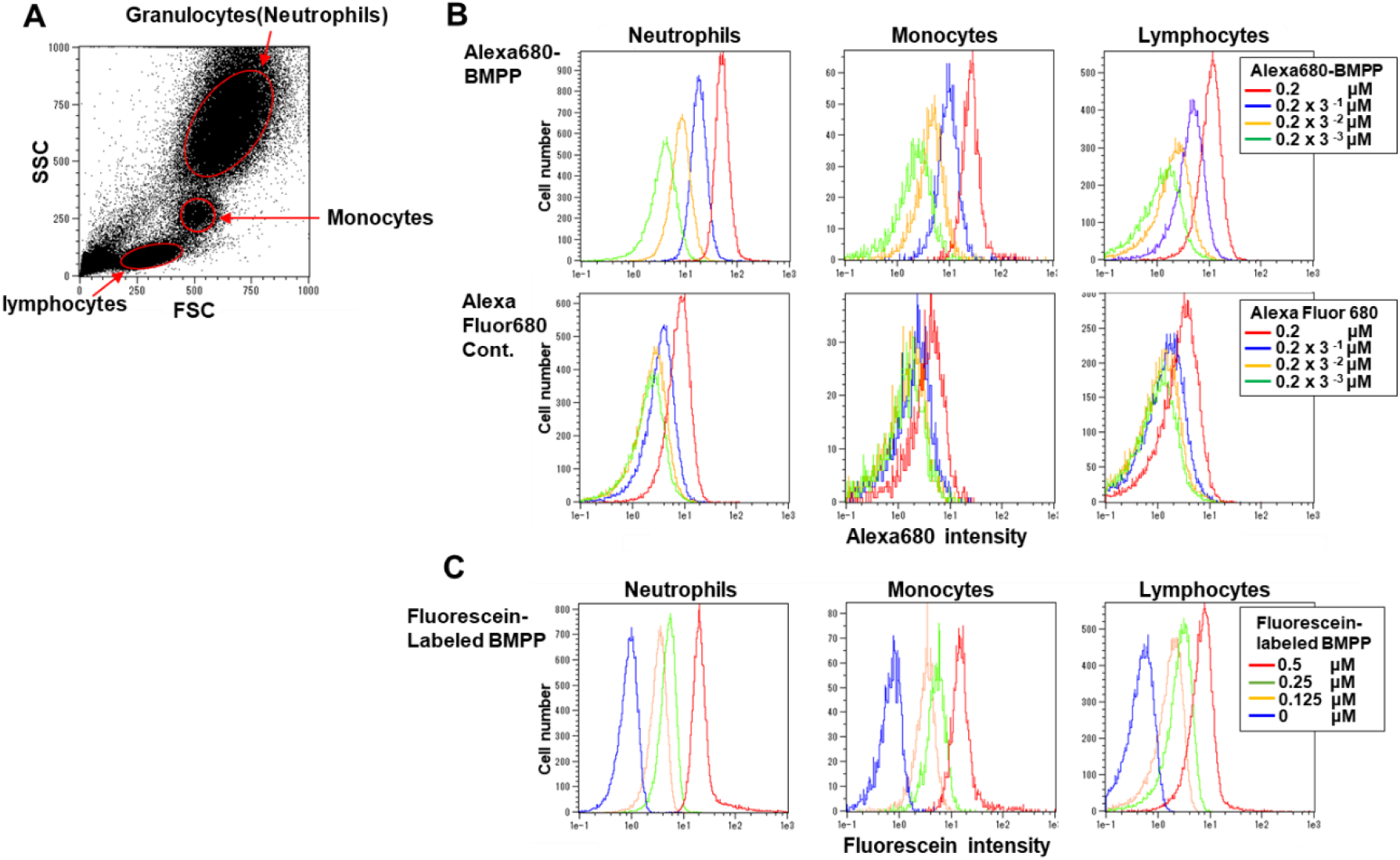
Dose-dependent specific incorporation of Alexa680-BMPP into white blood cells derived from human peripheral blood. (A) Dot plot of forward scatter (FSC) versus side scatter (SSC) showing human leukocyte subsets (granulocytes, monocytes, and lymphocytes). (B) Histograms showing the fluorescence intensities measured for blood cells treated with increasing doses (0.2 × 3^-3^ to 0.2 × 3^0^ μM) of Alexa680-BMPP (upper panels) or Alexa fluor 680 as a control (lower panels). (C) Histograms showing the fluorescence intensities measured for blood cells treated with increasing doses (0 to 0.5 μM) of fluorescein labeled BMPP. Concentration of fluorescently labeled BMPP are shown in each graph. Cell population names are indicated at the top of each graph. Note that the granulocyte gate delineated in (A) is considered to represent the neutrophil population.

### 3.2 White blood cells derived from a patient with CD36 deficiency incorporated Alexa680-BMPP

The uptake of ^123^I-BMIPP into cardiomyocytes occurs via a CD36 transporter that plays a crucial role in the uptake of LCFAs, reflecting the dynamics of LCFAs [13,14]. First, we assumed that WBCs could import Alexa680-BMPP via CD36 in the same manner as cardiomyocytes. Therefore, we prepared peripheral blood from a CD36-deficient patient and performed FACS analysis using Alexa680-BMPP. Surprisingly, the results showed that Alexa680-BMPP was incorporated into WBCs in a dose-dependent manner at the same level as the control (Fig. 2). This result suggests that WBCs have a unique pathway for LCFA export independent of CD36.

**Figure 2.**
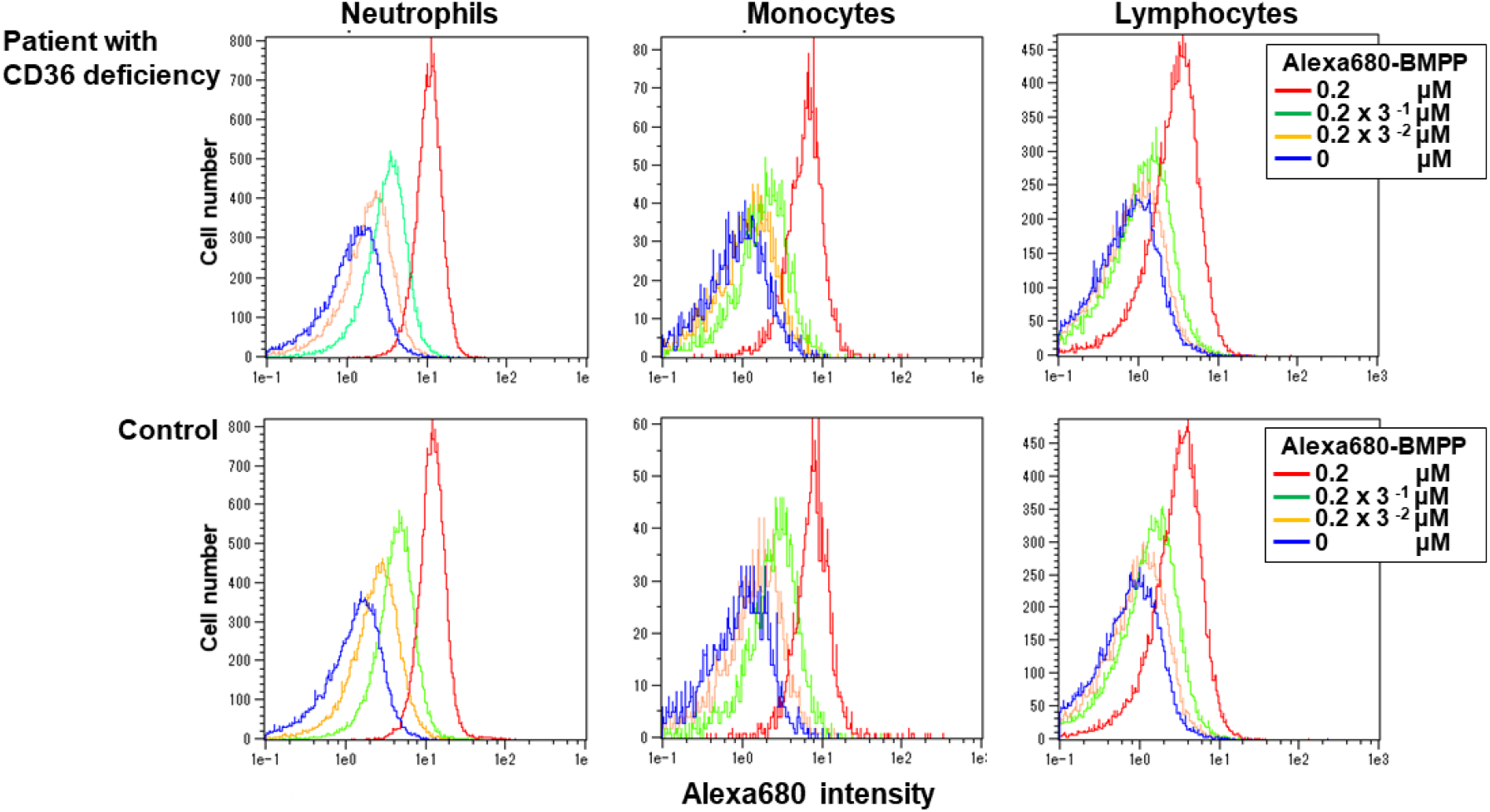
Incorporation of Alexa680-BMPP into white blood cells derived from a patient with CD36 deficiency. Histograms showing the fluorescence intensities of Alexa680-BMPP-incorporated blood cells derived from a patient with CD36 deficiency and a healthy volunteer (control). Upper and lower panels represent the CD36-deficient white blood cells and control, respectively. Concentrations of Alexa680-BMPP are shown in each graph. Cell population names are indicated at the top of each graph.

### 3.3 Export of Alexa680-BMPP was detectable following incorporation into white blood cells

To develop a fluorescent BMPP as an LCFA probe for TG metabolism of cells *ex vivo*, we examined whether the attenuation of Alexa680-BMPP fluorescence intensity in WBCs was detectable by FACS analysis after incorporation into cells. To this end, we employed a method in which Alexa680-BMPP was added to WBC fractions without erythrocytes to accurately investigate the export of Alexa680-BMPP after incorporation (section 2.4). The WBC fractions were treated with Alexa680-BMPP and then subjected to FACS analysis either immediately after treatment or after incubation in non-Alexa680-BMPP medium for 10 min to 2 h prior to FACS analysis (Fig. 3). Immediately after Alexa680-BMPP treatment, neutrophils, monocytes, and lymphocytes showed Alexa680 fluorescence intensities in line with the results described above (Fig. 3, 0 min). When cells were incubated in non-Alexa680-BMPP medium following Alexa680-BMPP treatment, the Alexa680 fluorescence intensities of neutrophils and lymphocytes were clearly attenuated after 10 min and did not differ significantly from those observed after further incubation (up to 120 min) (Fig. 3). Therefore, export of the fluorescent probe might have occurred rapidly. These results suggest that Alexa680-BMPP is a potent LCFA probe for the study of TG metabolism and LCFA import into cells. Hereafter, we focused on neutrophils to investigate the export of Alexa680-BMPP. Neutrophils were used in subsequent studies for several reasons. First, neutrophils showed dose-dependent incorporation of fluorescent BMPP in a relatively articulate manner. Second, neutrophils from patients with ATGL deficiency accumulate excessive neutral lipid vacuoles [15-17], suggesting that they substantially import FAs as an energy source. Third, the number of monocytes was remarkably decreased by the incubation. The reason why monocytes decreased after the incubation period is still unknown, although, it is likely that monocytes are vulnerable to damage during the experimental process of WBC fractions.

**Figure 3.**
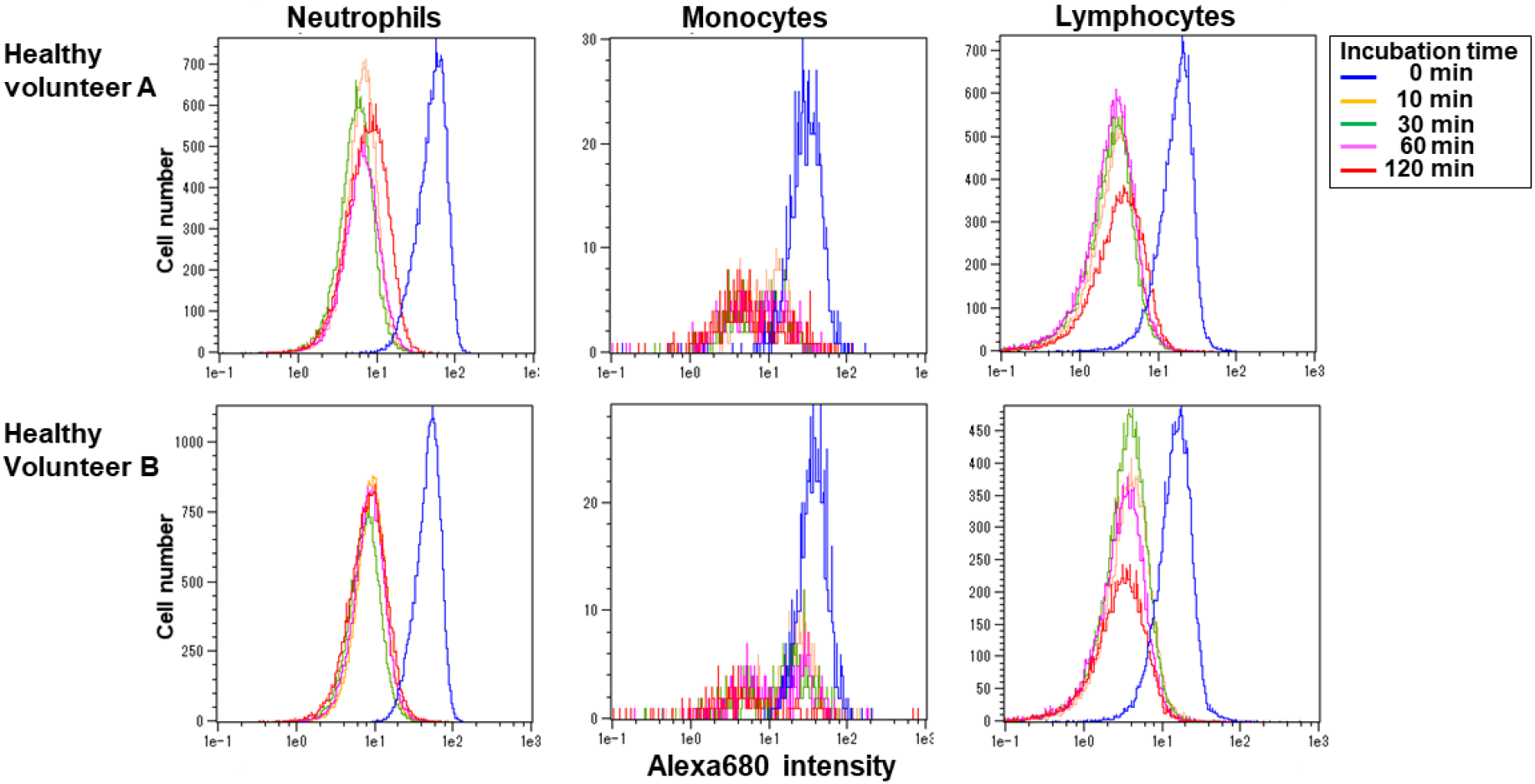
Alexa680-BMPP export from white blood cells. Histograms showing the fluorescence intensities of white blood cell fractions prepared from two healthy volunteers (Sample A and Sample B) that were treated with 0.75 μM Alexa680-BMPP. FACS analysis was performed after various incubation periods. Incubation times are shown in the Figure. Note that monocytes were largely lost after incubation (middle panels in both rows).

### 3.4 Decreased Alexa680-BMPP export was detectable in neutrophils derived from a patient with primary TGCV

Using WBC fractions derived from a patient with primary TGCV in which ATGL was deficient, we performed an *ex vivo* incorporation and export assay of Alexa680-BMPP using FACS. Results confirmed that the cells of a patient with primary TGCV showed a significantly decreased attenuation of Alexa680-BMPP fluorescence intensity compared to that of control cells after incubation in non-Alexa680-BMPP medium (Fig. 4), suggesting that patient-derived neutrophils had decreased activity for Alexa680-BMPP export. Therefore, it is highly likely that Alexa680-BMPP can be exported from cells mainly via ATGL-dependent lipolysis following its incorporation into the TG pool because ATGL deficiency caused a lower degree of fluorescence intensity attenuation. In addition, lymphocytes from the patient also showed decreased export activity, similar to that of neutrophils (Fig. 4). These results suggest that Alexa680-BMPP is a useful probe for the detection of TG metabolic deficiency in the WBCs of the patients.

**Figure 4.**
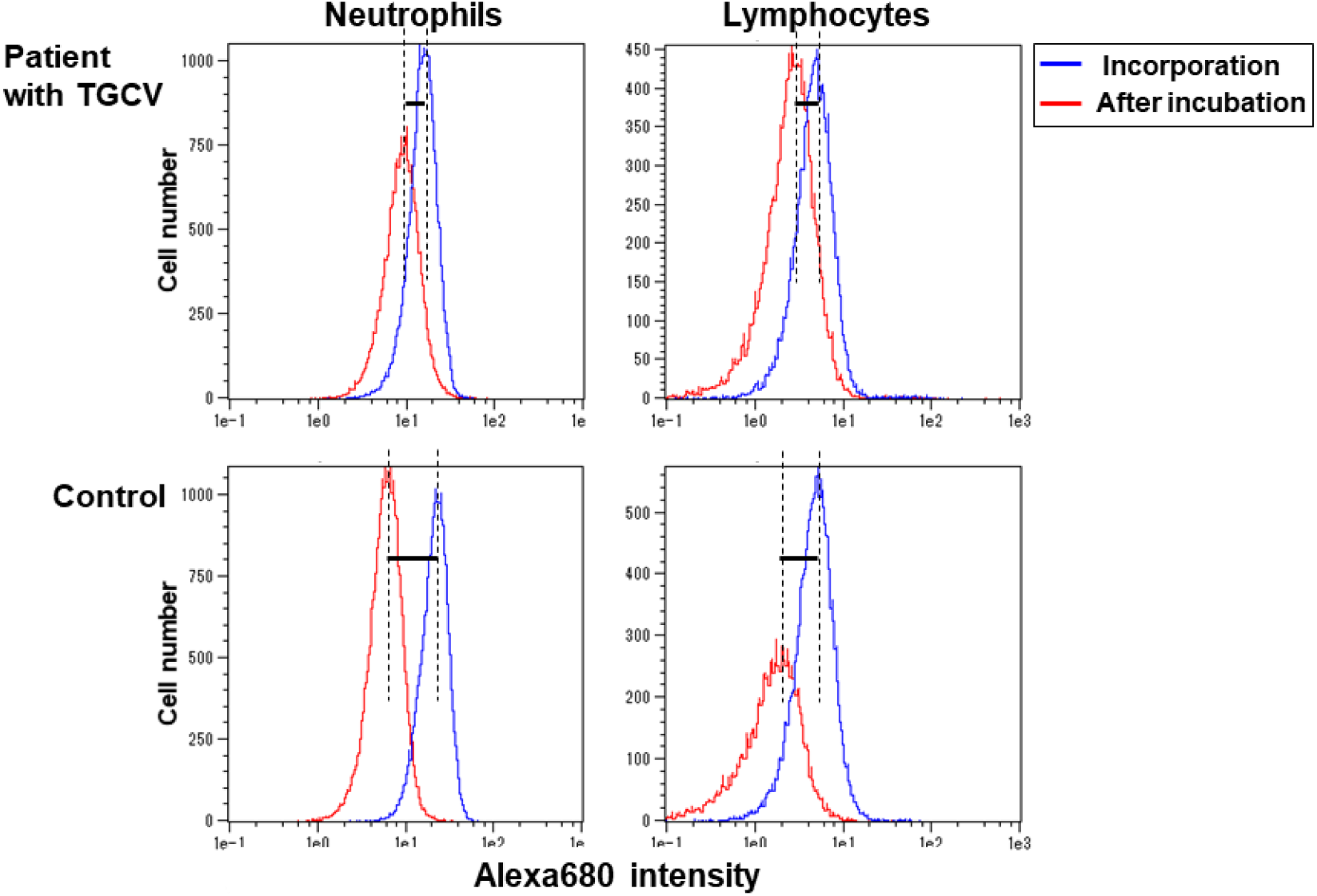
Decreased export of Alexa680-BMPP from neutrophils derived from a patient with primary TGCV. Histograms showing the fluorescence intensities of white blood cell fractions derived from a patient with primary TGCV (upper panels) and a healthy volunteer (control; lower panels) following an Alexa680-BMPP export assay. Blue and red colors indicate measurements taken before (immediately after Alexa680-BMPP incorporation) and after incubation (120 min), respectively. Dotted lines represent the positions of each peak, and solid horizontal lines between dotted lines represent the degree of export. Cell population names are indicated at the top of each graph.

### 3.5 Decrease of Alexa680-BMPP export in the cells of a patient with primary TGCV was ameliorated by treatment with tricaprin

We developed capsules containing tricaprin and conducted specific treatment for patients with primary TGCV [5]. Therefore, we compared the Alexa680-BMPP export activities of neutrophils derived from a patient before and after tricaprin treatment. The results showed a lower degree of attenuation of fluorescence intensity caused by ATGL deficiency and reverted to the same degree as that in the control cells after tricaprin treatment (Fig. 5). This likely reflects that the Alexa680-BMPP export activity in the patient’s cells was ameliorated by tricaprin treatment. Thus, the incorporation and export assay using Alexa680-BMPP may be able to detect fluctuations in TG metabolism, which reflects patient conditions.

**Figure 5.**
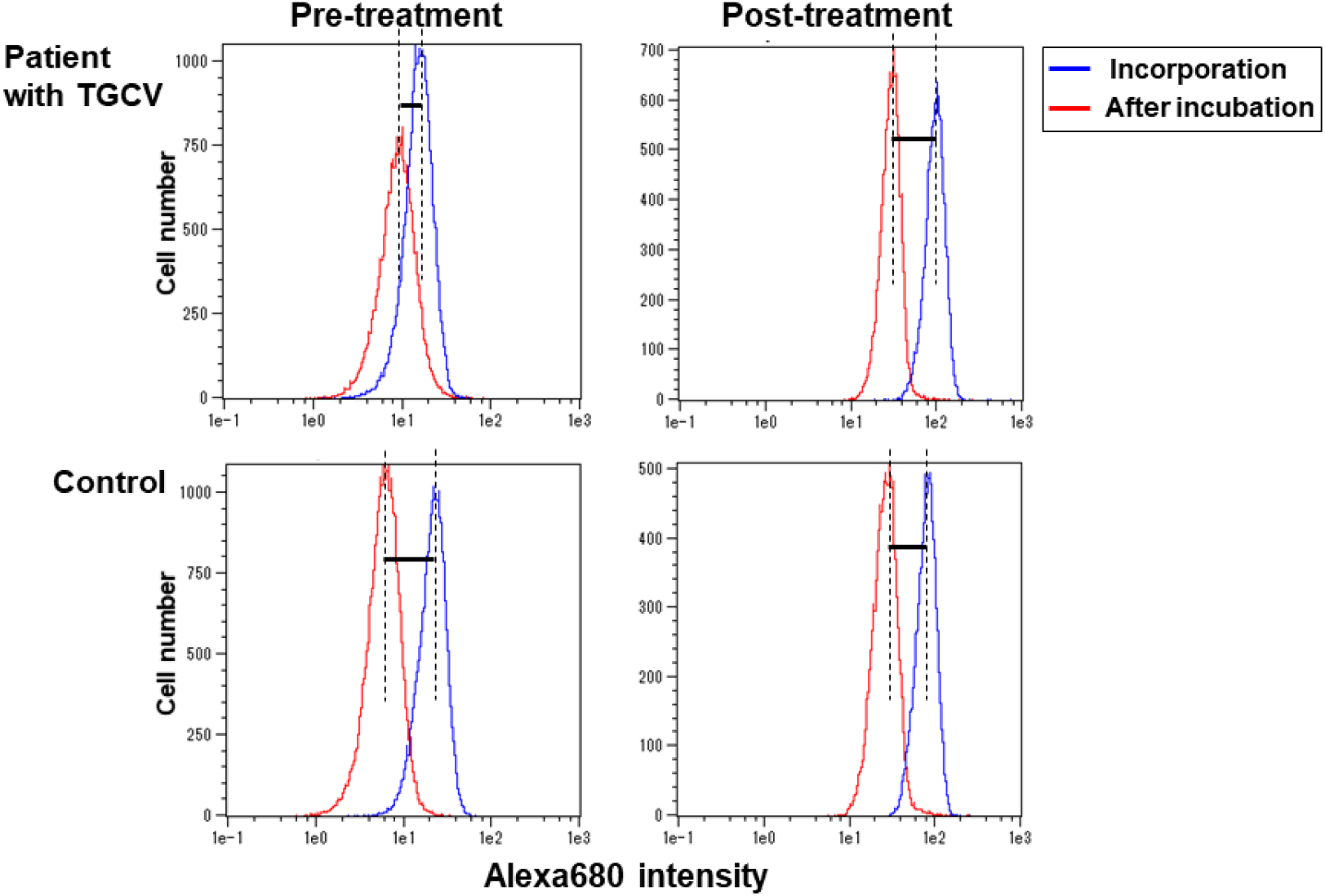
Comparison of Alexa680-BMPP export activity in a patient with primary TGCV’s neutrophils between post and pre-treatment with tricaprin. Histograms showing the Alexa680-BMPP export activity of neutrophils derived from a patient with primary TGCV after treatment with tricaprin (upper-right panel) compared to a healthy patient as the control (lower-right panel). The left panels show export activity before tricaprin treatment. The figure was delineated using the same format as that described in Figure 4. Note that the left panels, which represent the difference between the neutrophils of a patient with pre-treatment and in a control, are identical to those in Figure 4 (left panels).

## 4. Discussion

Given that ^123^I-BMIPP is commonly used as an LCFA-mimicking tracer in nuclear cardiology, we aimed to utilize an LCFA probe with the basic structure of BMIPP for *ex vivo* testing of intracellular lipase-dependent TG metabolism in cardiac diseases. The most distinctive feature of ^123^I-BMIPP is its methyl group at the β-3 position that inhibits direct mitochondrial β-oxidation and is retained in cardiomyocytes for a relatively long period of time [18,19]. This feature is convenient for comparing the lipase-dependent TG metabolism in cardiomyocytes. We previously performed myocardial scintigraphy with ^123^I-BMIPP in patients with TGCV who showed excessive accumulation of TG in the myocardium. Scintigraphy successfully detected prolonged retention of ^123^I-BMIPP in the heart and has, therefore, been established as a test for TGCV [5-7]. In the present study, using the advantages of the BMIPP basic structure, we utilized an Alexa680-labeled BMPP to detect LCFA metabolism in WBCs, particularly TG metabolism. To the best of our knowledge, this is the first study to use the fluorescently labeled BMIPP skeleton for examining the TG metabolism of cells derived from human samples together with FACS analysis.

In this study, we demonstrated that WBCs were able to incorporate fluorescent BMPP (Fig.1). Studies have reported that the uptake of ^123^I-BMIPP into cardiomyocytes is significantly impaired in CD36-deficient humans and CD36-knockout mice, suggesting that CD36 plays a central role [20-22]. First, we assumed that Alexa680-BMPP was imported into WBCs in a CD36-dependent manner, similar to ^123^I-BMIPP for cardiomyocytes. However, WBCs derived from a patient with CD36 deficiency could incorporate Alexa680-BMPP. Thus, WBCs are highly likely to have additional pathways to import Alexa680-BMPP independent of CD36, although the exact mechanism of import is still unknown.

^123^I-BMIPP is a powerful tool but it has disadvantages, such as the need for specific facilities and the burden of patients receiving the diagnostic procedure. Therefore, our goal is to develop a novel diagnostic method for TGCV that utilizes the BMIPP basic structure, but without a radioisotope and large equipment. To this end, we invented the concept of an *ex vivo* Alexa680-BMPP export assay using WBCs and FACS. We selected WBCs as an object for fluorescent BMPP for the *ex vivo* TG metabolism test for the following reasons. Circulating WBCs in the stationary state are known to utilize FAs as an energy source [1,2], which is consistent with the fact that FA oxidation is the predominant energy source for cardiomyocytes [23,24]. Furthermore, isolated leukocytes have been used for the direct detection of FA oxidation defects by radioisotope-labeled FAs, suggesting that leukocytes are suitable for *ex vivo* diagnosis of FA oxidation disorders [25,26]. In addition, circulatory neutrophils from patients with ATGL deficiency exhibit neutral lipid–containing vacuoles in the cytoplasm known as Jordans’ anomaly [15-17], suggesting that neutrophils aggressively use FAs as an energy source and are susceptible to TG metabolism deficiency. We successfully demonstrated that neutrophils from a patient with primary TGCV had lower Alexa680-BMPP export activity than those of the control, and this was ameliorated by tricaprin treatment, suggesting that the Alexa680-BMPP export assay is a possible method for detecting TG metabolism defects *ex vivo*.

However, this study had certain limitations. First, the method used before FACS analysis is extremely time-consuming. Thus, a strict time course for short periods cannot be observed given the fast transport of FAs into and out of cells. Second, TG metabolism deficiency diseases, including primary TGCV, are ultimately rare [5] and, therefore, samples can be hard to find. Hence, it was difficult to ascertain significant differences in the results of the Alexa680-BMPP export assay. These drawbacks need to be addressed in future studies. However, in this report, we demonstrated the possibility of using fluorescent BMPP in an *ex vivo* test for the diagnosis of primary TGCV.

In conclusion, this study presents a novel WBC TG metabolism assay using a fluorescently labeled LCFA analog probe with a BMIPP-based structure (Alexa680-BMPP). This assay is based on the *ex vivo* detection of Alexa680-BMPP incorporation and export by neutrophils using FACS analysis. Our findings show that Alexa680-BMPP export activity is decreased in neutrophils from a patient with primary TGCV compared to that in a healthy control. Furthermore, clear amelioration in Alexa680-BMPP export activity was confirmed in neutrophils from the same patient after tricaprin treatment. Taken together, this probe has great potential for *ex vivo* testing of TG metabolism in WBCs derived from patients with lipid metabolism deficiencies, including coronary artery disease, metabolic syndrome, and primary TGCV.

## Authors’ contributions

YH designed the experimental protocol, performed experiments, and wrote drafted the manuscript. KH designed the study and coordination and wrote the manuscript. The authors have read and approved the final manuscript.

## Acknowledgements

We thank Chikako Hashimoto for collecting the patients’ peripheral blood. We are grateful to Professor Kenji Mode (Hokkaido University) and Professor Takashi Jin (Center for Biosystems Dynamics Research, RIKEN) for providing fluorescently labeled BMPP. We also thank Professor Hirokazu Kashiwagi (Osaka University) for his useful suggestions and helpful comments. We thank all participants.

This study was partially supported by research grants for rare and intractable diseases from the Japan Agency of Medical Research and Development (AMED) (Grant No. 17ek0109092h0003) and the Ministry of Health, Labour, and Welfare of Japan(24FC1007).

## Ethics

These studies conformed to the principles outlined in the 1964 Declaration of Helsinki and were approved by the Osaka University Hospital Ethics Committee (Approval Nos. 09122 and 14207). Written informed consent was obtained from all patients treated with tricaprin.

